# Quantifying Physiological Synchrony Using Windowed Cross-Correlation Analysis: Statistical and Theoretical Considerations

**DOI:** 10.1101/2020.08.27.269746

**Authors:** F. Behrens, F. Diana, R. G. Moulder, S. M. Boker, M. E. Kret

**Affiliations:** Leiden University, Institute of Psychology, Cognitive Psychology Unit, Wassenaarseweg 52, Leiden 2333 AK, The Netherlands; Department of Social, Health and Organizational Psychology, Utrecht University, Heidelberglaan 1, 3584 CS Utrecht, The Netherlands; Department of Psychology, University of Virginia, Charlottesville, VA 22903, USA

**Author notes:** **Corresponding Author:** M. E. Kret, Leiden University, Institute of Psychology, Cognitive Psychology Unit, Wassenaarseweg 52, Leiden 2333 AK, The Netherlands;.

**Keywords:** Windowed Cross-Correlation analysis, interpersonal synchrony, parameter optimization, surrogate data, time series analysis

## Abstract

Interpersonal synchrony is a widely studied phenomenon. A great challenge is to statistically capture the dynamics of social interactions with fluctuating levels of synchrony and varying delays between responses of individuals. Windowed Cross-Correlation analysis accounts for both characteristics by segmenting the time series into smaller windows and shifting the segments of two interacting individuals away from each other up to a maximum lag. Despite evidence showing that these parameters affect the estimated synchrony level, there is a lack of guidelines on which parameter configurations to use. The current study aimed to close this knowledge gap by comparing the effect of different parameter configurations on two outcome criteria: i) the ability to distinguish synchrony from pseudosynchrony by means of surrogate data analyses and ii) the sensitivity to detect change in synchrony as measured by the difference between two within-subject conditions. Focusing on physiological synchrony, we performed these analyses on heartrate, skin conductance level, pupil size, and facial expressions data. Results revealed that a range of parameters were able to discriminate synchrony from pseudosynchrony. Window size was more influential than the maximum lag with smaller window sizes showing better discrimination. No clear patterns emerged for the second criterion. Integrating the statistical findings and theoretical considerations regarding the physiological characteristics and biological boundaries of the signals, we provide recommendations for optimizing the parameter settings to the signal of interest.

## Introduction

During social interactions, humans tend to synchronize on different levels. They mimic motor movements, such as postures (Tia et al., 2011), facial expressions (Altmann et al., 2021; Riehle et al., 2017), scratching and yawning (Diana et al., 2023; Schut et al., 2015),and motion activity (Altmann et al., 2020) and align their level of physiological arousal (Feldman et al., 2011; Levenson & Gottman, 1983; Prochazkova et al., 2018). Although this synchrony comes naturally and without effort, it is a great challenge for social scientists to measure it statistically, particularly for physiological synchrony, where signals are continuous and constantly fluctuating. This challenge is especially pronounced in naturalistic interactions, which are shaped by a continuous back-and-forth of signals and responses between individuals, with varying time delays and fluctuating levels of alignment that are difficult to statistically capture. To address this challenge, the current paper builds on previous work on movement synchrony (Boker et al., 2002) to propose Windowed Cross-Correlation (WCC) as a solution to capture the dynamic changes in heartrate, skin conductance level, facial expression, and pupil size (Boker et al., 2002). We first describe the WCC method and the challenge of selecting appropriate parameter configurations. We then present an empirical comparison of parameter settings across two criteria, the ability to distinguish synchrony from pseudosynchrony, and the sensitivity to detect changes in synchrony, tested on physiological data from a face-to-face storytelling paradigm. Based on these comparisons, our aim is to provide recommendations to help researchers make well-informed decisions in applying the WCC analysis.

Synchrony is a multifaceted phenomenon evident on the behavioral, physiological, and neural level. Not surprising then, the causes and consequences of synchrony have been studied in a broad range of contexts investigating the dynamic nature of social interactions from clinical (Galazka et al., 2019; Wehebrink et al., 2018), developmental (de Klerk et al., 2018; Shih et al., 2019), evolutionary (Diana & Kret, 2025;Palagi et al., 2009), neural (Hasson et al., 2004; Prochazkova et al., 2018), social (Behrens et al., 2020; Tarr et al., 2016; Diana et al., 2025), and cognitive (Kret et al., 2015; Kret & De Dreu, 2017) perspectives. Crucially, many of the interpersonal processes influenced by synchrony, such as therapeutic interactions (Ramseyer & Tschacher, 2011), mother-infant exchanges (Feldman et al., 2011), and trust decisions (Diana et al., 2025), are inherently lagged in nature, where one person’s signal triggers a delayed response in the other and lead-follow dynamics shift across the course of an interaction.

A variety of methods have been proposed in previous literature to quantify synchrony including correlations, regressions, structural equation models and recurrence quantification analysis. These approaches differ in their assumptions, their operationalization of synchrony, and the type of synchrony they measure (for a review, see (Gates & Liu, 2016; McAssey et al., 2013; Schoenherr et al., 2018; Thorson et al., 2017). For continuous physiological time-series, which are the focus of this article, an ideal method needs to fulfill three criteria: First, it should capture both simultaneous (in-synch) responses, e.g., where two individuals react simultaneously to an external event, and lagged responses, where one individual responds to another individual or at a different pace. Second, it should allow for changes in the level of synchrony across the interaction, reflecting moments of stronger and weaker coupling. Third, rather than categorizing intervals as synchronous or not (as in Altmann, 2011), it should provide a continuous estimate of synchrony strength. This is particularly important for physiological signals, which are constantly fluctuating and rarely settle into discrete states that can cleanly be labeled as synchronous or non-synchronous. For instance, two individuals’ heartrates may partially overlap in their fluctuations for a period without ever reaching a clear threshold of synchrony, yet this partial coupling may still carry meaningful information about their interaction. While reducing synchrony to a binary classification is a valuable approach for movement synchrony, where people can either move or not, it may not be optimal for physiological signals. A method that fulfills these different criteria is Windowed Cross-Correlation analysis, the focus of the current study (Boker et al., 2002).

Windowed Cross-Correlation (WCC) analysis offers a neat method to account for dynamic changes in synchrony (Boker et al., 2002). This is achieved by extending a classical cross-correlation estimate by two aspects: **windows** and **lags**. Rather than calculating a correlation coefficient over the whole time series, the signals are broken into smaller overlapping segments or windows. The strength of alignment to be estimated separately for each window, thus allowing synchrony to vary from stronger to weaker across the interaction. Introducing the lag is useful to account for the natural back-and-forth dynamics of social interaction: within each window, segments are shifted relative to one another up to a maximum lag, capturing the lead-follow dynamics between individuals (e.g., Person A responds to Person B and a moment later the pattern is reversed). Despite these advantages, the method requires researchers to specify parameters that tailor the analysis to the signal of interest. While Boker et al. (2002) provided guidance on these parameters for motor movement data, no guidelines exist for physiological measures. The current paper aims to close this gap.

### WCC parameters

The WCC analysis requires the specification of **four parameters** that tailors the method to the signal of interest: window size, maximum lag, window increment, and lag increment (see Figure 1). Carefully choosing the right parameter settings is crucial, because these settings can substantially affect the outcome of the WCC (Schoenherr et al., 2018).

**Figure 1.**
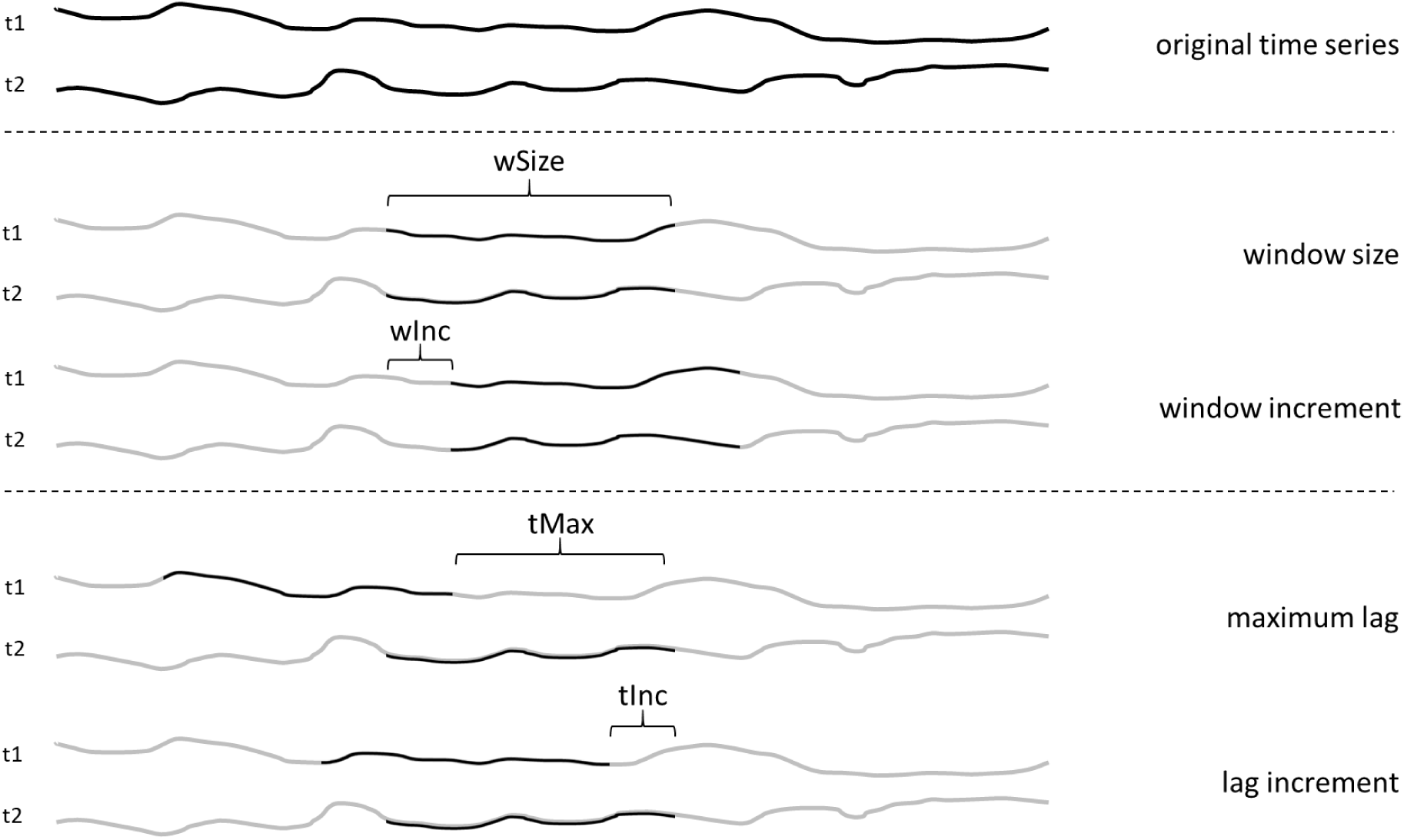
Schematic outline of the four parameters that are specified in the WCC analysis: window size (wSize), window increment (wInc), maximum lag (tMax), and lag increment (tInc). The abbreviations tMax and tInc originate from using ‘tau’ (τ) to refer to the lags in the cross-correlation equation. ^i^

#### Window size

The window size determines the number of samples in each sliding window across the time series. It should be small enough to be sensitive to changes in the synchrony strength, in the lead-follow relationship between individuals, and to satisfy the assumption of local (Boker et al., 2002). Disregarding fluctuations within a large window might undermine the strength of association at certain moments. However, the window should be large enough (i.e., have enough data points) to provide reliable correlation estimates. While 50-70 values have been proposed as sufficient (Cappella, 1996), more recent work performing Monte-Carlo simulations recommends 65 to 250 values, depending on the strength of the correlation (Schönbrodt & Perugini, 2013). Given the high sampling rates incorporated in many psychophysiological measurement devices, this range should be fairly easy to accomplish, if the window size is not overly small. The biological nature of the signal also matters: slower signals such as skin conductance require longer windows than faster signals such as facial expressions. Decisions on the window size should therefore balance both statistical and theoretical considerations.

#### Maximum Lag

The maximum lag indicates how far (i.e., the maximum number of samples) one window is shifted relative to the other, and consequently defines the maximum delay within which two responses are still considered reactions to one another. For example, if the maximum lag is three seconds and Person A smiles two seconds after Person B, this would be captured; if that smile occurs four seconds later, it would not be considered a response anymore. If the maximum lag is too long, synchrony might be attributed to unrelated events; if too small, meaningful delayed responses may be missed. Prior work suggests that the maximum lag matters: for instance, skin conductance responses within but not beyond seven seconds have been shown to correlate with empathetic counselor-client relationships (Robinson et al., 1982). However, this study did not directly compare whether shorter latencies predicted the relationship better than longer ones. Furthermore, the broad categorization of latencies (responses between 0 and 7 seconds compared to responses between 7 and 40 seconds) does not allow for fine-grained conclusions about which maximum lag is optimal. Thus, although this provides an indication that the maximum lag indeed matters, a systematic comparison of different maximum lags is needed to make well-informed decisions on this parameter.

#### Window Increment and Lag Increment

The window increment determines the size of the steps (i.e., the number of samples) when moving from one window segment to the next, regulating the temporal resolution of the analysis. When the increment equals one, then the window is moved by one data point; when it is the same size as the window size or greater, adjacent windows are non-overlapping. Similarly, the fourth parameter, the lag increment, indicates how big the steps are between time lags. For both parameters, smaller increments yield better resolution but at the cost of increased computational time. Crucially, however, estimates will eventually stabilize as increments decrease, making the information that can be added not worth of the increased computational time. Comparing it to sampling rates, if one aims to measure heartrate changes, a sampling rate of 1000 Hz gives a smooth signal. Increasing the sampling rate to 2000 Hz adds little information because the heartrate does not change this fast and so very similar heartrate signals using both sampling rates. Similarly, increasing the resolution of the increment of the moving windows and lags will eventually stabilize around a correlation estimate. Naturally, the lower bound of both increments parameters will be determined by the sampling rate of the device. Setting these parameters is therefore a matter of balancing resolution against computational efficiency.

### Criteria to determine the best parameters

To determine the best parameter configurations, we used two criteria. The first criterion was the ability to discriminate synchrony from pseudosynchrony, defined as the spurious synchrony that arises between individuals not engaged in actual information exchange (Moulder et al., 2018). Such spurious synchrony exists because physiological signals are constrained in their patterns: heartrate, for instance, fluctuates within a certain range regardless of social context, creating baseline similarities between any two individuals’ signals, even more so if individuals are engaged in similar activities (i.e., participating in a study with the same procedure across dyads). However, the changes occurring during spurious synchrony stay in a certain range causing recursiveness and commonality within and between heartrate measures. Consequently, the null hypothesis for synchrony testing is not zero but some fundamental baseline value, necessitating an appropriate comparison. It is therefore necessary to find an appropriate comparison between the level of synchrony of individuals engaging in an interaction and the level of synchrony that occurs due to the nature of the signals. One way to account for pseudosynchrony is to perform a surrogate data analysis (Moulder et al., 2018). The idea is that the original or true time-series is compared to the same time-series where synchrony is destroyed while keeping all other properties constant. Specifically, synchrony levels from true interacting dyads are compared to those from newly generated dyads of participants who never actually interacted. To generate these dyads, the time-series from each participant is coupled with every other participant, testing whether being in an interaction adds something over and beyond what is expected from shared situational and biological factors alone.

The second criterion was the sensitivity to detect changes in synchrony in response to experimental manipulations, a key requirement for studying the underlying mechanisms, boundary conditions and individual differences in synchrony. For example, previous studies, we observed that physiological synchrony affected behavioral outcomes, but only when participants could see each other (our manipulation) (Behrens et al., 2020; Diana et al., 2025). Another study investigated the effect of emotional salience during storytelling on pupil mimicry and showed that physiological coupling between the speaker and the listener was stronger during emotionally intense moments compared to less salient moments (Kang & Wheatley, 2017). Storytelling is a particularly interesting setting. It is a uniquely human and universal activity creating social bonds between people (Smith et al., 2017), and it is also an inherently lagged activity where one participants listens and the other tells the story.

Based on the lagged nature of story-telling and the effect of face-to-face contact on physiological synchrony, in the current study two individuals engaged in face-to-face storytelling. Participants also completed baseline measures of eye contact in silence. In line with the findings by Kang and Wheatley (2017), we expect higher levels of synchrony when people engage in storytelling compared to the baseline measure. These two conditions are ideal to test whether the WCC analysis we use to compute synchrony will be sensitive enough to detect *changes* in synchrony between the two (within-subject) conditions. The aim of the current study is to determine the best parameter configurations for the Windowed Cross-Correlation (WCC) analysis applied to different common physiological measures. In line with the definition provided by Altman (2011), we operationalize synchrony as a temporary linear relationship between two time series, focusing on time series describing physiological changes over time in two individuals. We tested the two criteria outlined above on data from dyadic interactions where two individuals told each other four stories. During the interaction, their heartrate, skin conductance level, pupil size, and contractions of the left zygomaticus major (muscle associated with smiling) were measured. We chose these four measures as they have all been shown to synchronize between individuals, include both sympathetic and parasympathetic responses, and are prominent measures in the literature (for recent reviews, see Palumbo et al., 2017; Prochazkova & Kret, 2017; Diana & Kret, 2025). For a range of window sizes and maximum lags that were tailored to each signal, we calculated a measure of distance for the comparison (i) between the true dyads and newly generated surrogate dyads, and (ii) between intervals of storytelling and baseline measures in the true dyads. The window and lag increments were not systematically compared, but were adjusted as a function of the window size and maximum lag, respectively. Based on the outcome of these comparisons, we provide recommendations on which parameter configurations are best for detecting synchrony and change in synchrony for the four physiological measures.

## Method

### Participants

In total, 34 same-sex dyads participated in the study, of which six dyads had to be excluded due to technical problems (dyads included in analysis: Female = 22 (78%), Male = 6 (22%); *M_age_* = 22.79; *SD_age_* = 3.23; Dutch = 17 (30%)). Participants were recruited via the Leiden University online recruitment system, flyers distributed around the university building, and through personal contacts. In the latter case, the participant was tested by a researcher they did not know. The relationship between the two participants was not formally assessed, but verbal questioning confirmed that no dyad consisted of (close) friends. Individuals had normal or corrected-to-normal vision wearing contact lenses. Glasses were not compatible with the eye-tracking glasses worn during the experiment. The duration of the study was about one hour and participants received two course credits or 6€, and chocolate for compensation. The study was approved by the local Psychology Ethics Committee of Leiden University (CEP19- 0313/208).

### Design

The design of the study is outlined in Figure 2. The study consisted of two parts: (1) participants completed a breathing exercise where they were instructed to look at each other and synchronize their breathing; and (2) participants engaged in storytelling with each participant telling a neutral and a positive story while the other participant was listening. In total, participants told each other four stories, with Stories 1 (positive) & 3 (neutral) told by participant 1 while participant 2 was listening, and Stories 2 (positive) & 4 (neutral) being told by participant 2 while participant 1 was listening, with the starting stories order was counterbalanced between dyads. The breathing and storytelling parts were both preceded by a 2-min baseline measure where participants were instructed to relax and look at each other. After the second baseline measure and after each story, participants filled out the PANAS (Positive And Negative Affect Schedule – Watson, et al., 1988) to measure their current affect. Furthermore, they rated each story with regard to its valence and intensity on a scale from 0 to 10. The PANAS, the story ratings, and the rest of the questionnaires scores are beyond the scope of this study, but the descriptive statistics are provided in Table S1 (Supplementary Materials).

**Figure 2.**
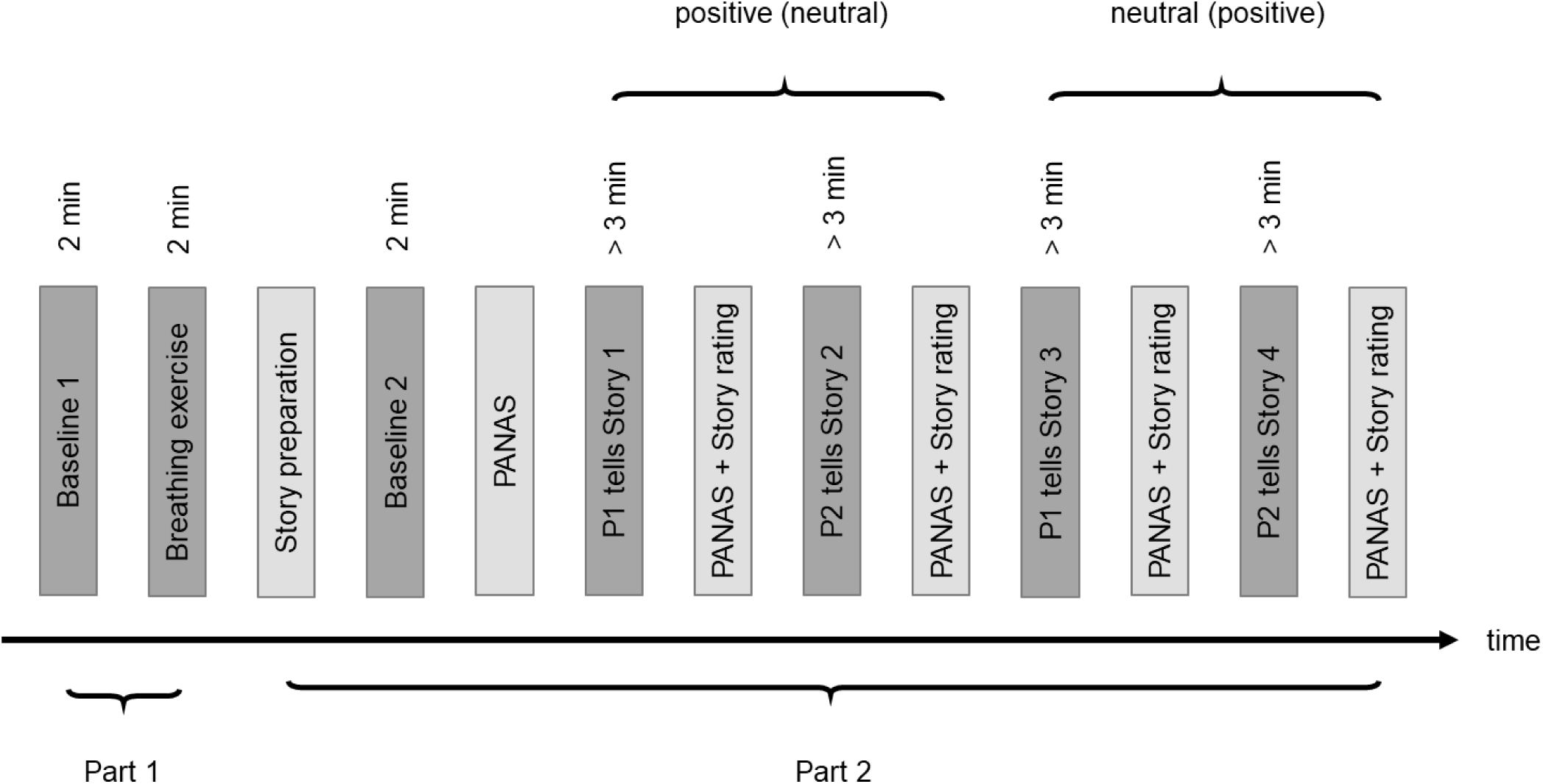
The time course of the study. The study was divided into two parts: breathing exercise (Part 1) and storytelling (Part 2). During the dark grey epochs, people interacted with each other; during light grey epochs, they prepared the storytelling and filled out questionnaires; P1/P2 = Participant 1 and 2; PANAS = Positive And Negative Affect Schedule; Story 1 & 2 and Story 3 & 4 were of the same valence (positive or neutral), the order of starting with the positive or neutral story was counterbalanced between dyads.

### Procedure

Upon arrival at the lab, participants were separated, received information about the study, and gave informed consent for participation. Afterwards, electrodes were attached to the face, fingers, and the torso as preparation for the measurement of EMG, EDA, and ECG activity, respectively. Specifically, two electrodes were attached to the non-dominant hand on the intermediate phalanges of the index and ring finger to measure skin conductance level; three electrodes were attached on the left and right side of the abdomen and on the thorax below the right collar bone to measure heartrate; and three electrodes were attached to the left face on the zygomaticus major and behind the ear to measure facial expressions. After the preparation, participants filled out the Interpersonal Reactivity Inventory (IRI – Davis, 1980) and the Five Facet Mindfulness Questionnaire (FFMQ; Baer, et al., 2006). Next, participants were seated on the same table, and wore the eye-tracking devices Tobii Pro Glasses 2, which were subsequently calibrated. Afterwards, the experimenters left the room and started the recordings of the physiological measures. Participants received pre-recorded instructions via speakers. The breathing, baseline and storytelling parts were completed as described in the Design section above and outlined in Figure 2. At the end, participants put all filled out papers in an envelope, and were debriefed and rewarded for their participation.

### Preprocessing of the physiological measures

The physiological measures were recorded with the MP160 BIOPAC data acquisition system at a sampling rate of 2000 Hz and were pre-processed offline with the PhysioData Toolbox (Sjak-Shie, 2017). Using the Toolbox’s standard preprocessing settings, the heartrate data were preprocessed applying a band-pass filter between 1Hz and 50Hz. R-peaks were detected and transformed to inter-beat intervals (IBI) and subsequently to heartrate (bpm) values. The skin conductance signal was low-pass filtered with a cut-off of 5Hz. The EMG signal was preprocessed with a low-pass FIR filter of 28Hz and a high-pass FIR filter of 500Hz and a Notch-filter of 50Hz. The rectified signal was subsequently smoothed with a Boxcar filter of 100ms. The pupil size data were preprocessed in multiple stages according to recommended guidelines described elsewhere (Kret & Sjak-Shie, 2018). After applying the filters, each signal was visually inspected. If missing or incorrect intervals were manually detected, the signals were linearly interpolated. Finally, all signals were down-sampled to 20Hz.

### Windowed Cross-Correlation analysis

Two challenges in analyzing physiological responses between two individuals include i) to statistically represent the dynamics of an interaction and ii) to quantify the associated patterns in that might vary in the strength of association and the timing of the responses. Windowed Cross-Correlation (WCC) analysis offers a method that addresses both challenges. Specifically, the two time-series are broken into smaller, overlapping windows before the correlation is estimated for each window. This way, the strength of association can vary between these windows accounting for the non-stationarity of the signals. The overlap between windows assures that strong synchronization that occurs at the edge of non-overlapping adjacent segments is not missed. Additionally, for each window, the two segments are lagged away from each other up to a maximum lag such that the segment of either participant 1 or participant 2 precedes the other participant’s segment in time, accounting for the (varying) delay between individuals’ responses. This generates a result matrix r with correlations for the different segments and time lags defined as

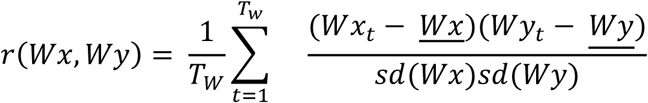

Where *T_w_* is the total amount of observations in each window *Wx* and *Wy* consisting of observations *Wx_t_* and *Wy_t_* where *t* Є {1, …, *T_w_*), *<u>Wx</u>* and *<u>Wy</u>* are the means of the observations in each window, and *sd*(*Wx*) and *sd*(*Wy*) the standard deviations of each window. In the result matrix, each row represents one window, while each column represents one lag. Because the first window needs to lag segments up to the maximum lag and because the window includes more than one data point, the number of rows is given by (*N* – *wSize* – *tMax*) / *wInc*. Dividing by *wInc* accounts for how many observations are skipped between one window and the next one. For example, if the window increment is one, then the number of rows of the result matrix will be equal to the number of observations of the time-series (after accounting for the window size and maximum lag as just described). But if the increment is 10, then the steps are bigger between the windows, reducing the number of segments needed to cover the whole time-series and therefore decreasing the number of rows in the result matrix. The number of columns in the result matrix is (*tMax* * 2) / *tInc* + 1 because the segments are shifted such that first Participant 1 and then Participant 2 precedes the other participant up to the maximum lag (i.e., twice the *tMax*). The *tInc* accounts for the size of the steps between two lags. The extra column (+1) represents the case where the lag is zero.

### Peak picking

Following the WCC analysis, Boker et al. (2002) developed the so-called peak-picking algorithm where the maximum correlation across different lags is determined for each window (i.e., the maximum correlation per row of the result matrix). The maximum correlation should be preceded and succeeded by lower correlation values. For example, if Participant 1 synchronizes with Participant 2 with a lag of 1 second, then the correlation should be highest (i.e., peaks) at that time lag and the correlation should be lower at both lag .5 and 1.5 seconds. This “peak” is implemented to ensure that individuals indeed react to one another. If both individuals did nothing, they both would show more or less flat lines in their physiological responses and the correlation between their signals would be high for all lags. Requiring a peak in the correlation across lags prevents such events from being termed “synchrony”. The peak picking algorithm outputs a matrix with the maximum (i.e., peak) correlation with its corresponding time lag for each window. In a last step, a summary statistic is computed by calculating the mean of the maximum correlations. This measure provides an indication of the overall level of synchrony between the two time-series.

### Choosing values for parameter configurations

To quickly recap, there are four parameters that need to be specified: window size, window increment, maximum lag, and lag increment. The window size (*wSize*) determines how long each window is, the window increment (*wInc*) indicates the size of the steps between two adjacent windows, the maximum lag (*tMax*) regulates how far the segments of the two time-series are shifted away from each other, and the lag increment (*tInc*) determines the size of the steps with which the segments are shifted.

To determine the range of values for the window size and maximum lag parameters, we employed a bottom-up approach. Fist, we run preliminary WCC analyses on the whole time-series (including all data of the study), and visually inspected the resulting matrix plots. Optimal parameter configurations produce sharp contrasts between regions of high and low synchrony, whereas suboptimal configurations yield a smoothed image with less contrast, making differences harder to detect (see Figure 3). For the maximum lag, we additionally checked whether peak correlations fell within the range of lags or outside the plot boundaries, which would indicate the lag range was too narrow. Second, we ensured the range of parameters included values previously used in the literature. Third, we set the minimum window size to 3 seconds, corresponding to at least 60 data points at our 20Hz sampling rate, in line with guidelines for reliable correlation estimation (Schoeneberger, 2016). For the sake of simplicity, the range of maximum lags was equal to the range of window sizes. The window size and maximum lag parameters chosen for each physiological measure are listed in Table 1. For the window and lag increment parameters, we used 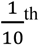 of the window size and the maximum lag, respectively.

**Figure 3.**
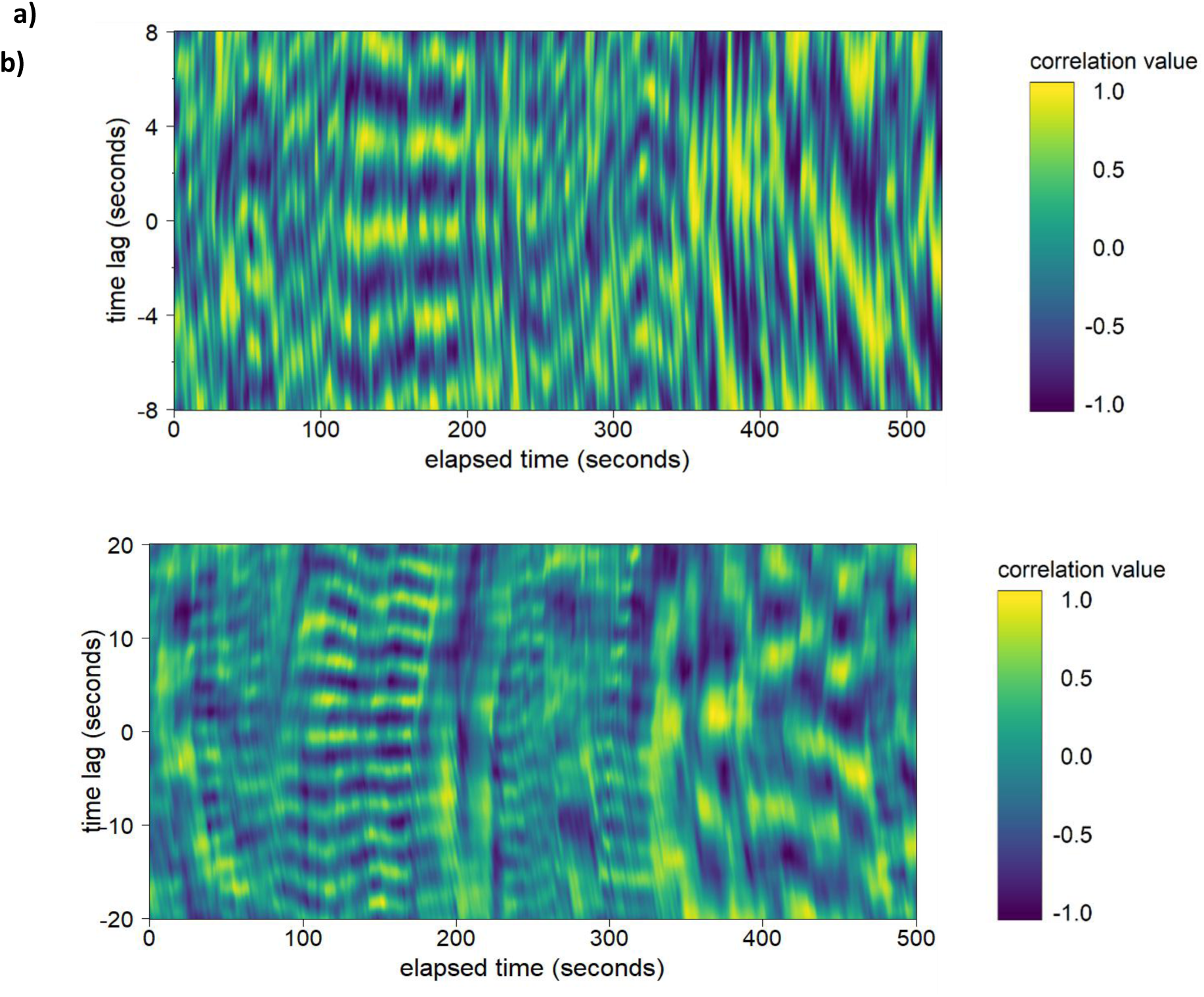
Examples of WCC analysis plots using heartrate data and a window size and a maximum lag of (a) 8 sec and (b) 20 sec, representing a optimal and suboptimal example of parameter settings, respectively. Between around 100 and 200 seconds, people engage in a breathing exercise where they breathe synchronously which is reflected in the steadily high correlations around the time lag of zero.

**Table 1:**
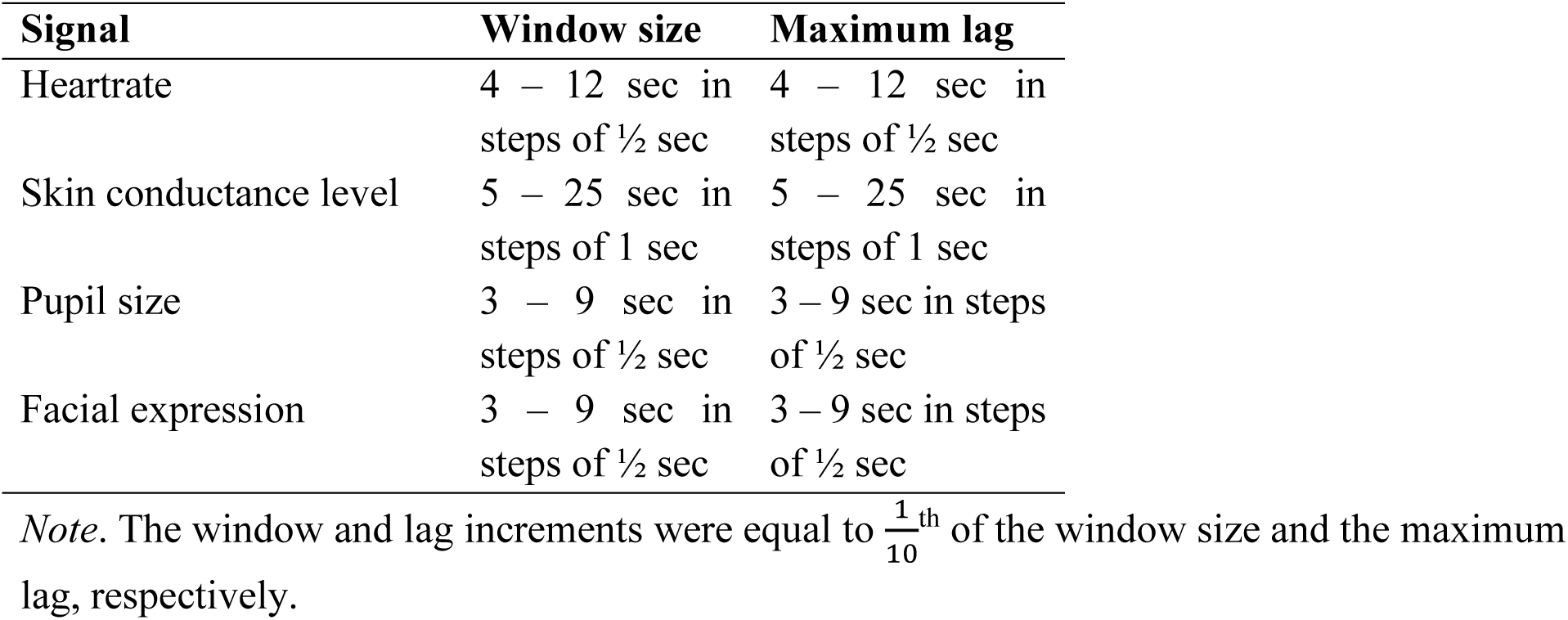
Window size and maximum lag parameters used for each physiological measure.

### Choosing the best parameter settings

We conducted the WCC and peak picking analysis for all combinations of the window size and maximum lag parameters with their corresponding increments as described in the previous section. For each parameter configuration, we calculated the mean peak correlation across window segments per dyad as the measure of synchrony. To determine the best parameter configurations for each physiological measure we used the two criteria outlined in the introduction: (i) the ability to discriminate synchrony from pseudosynchrony, and (ii) the ability to detect *change* in synchrony.

For the first criterion, we compared the true dyads (i.e., individuals who actually interacted during the experiment) with the surrogate dyads (i.e., all possible pairings of pairing individuals who did not interact). If the specific social interaction evoked synchrony above and beyond the synchrony shared situational factor, synchrony levels should be higher in the true compared to the surrogate dyads. As such, we computed the mean peak correlation for true and the surrogate dyads, testing whether specific parameter configurations were more sensitive in detecting the difference between synchrony (true dyads) and pseudosynchrony (surrogate dyads). Sensitivity to detect this difference was quantified using the t-statistic of an independent samples t-test between the mean peak correlations of the two groups, with a positive t-statistic indicating higher synchrony in the true dyads. Importantly, we used the t-statistic purely as a measure of distance between group means rather than for hypothesis testing, and therefore interpret results in relative rather than absolute terms and do not draw any conclusions about whether the differences reveal significant results or not. The analysis was conducted on data from the first baseline measure and replicated on the second baseline measure to assess robustness.

For the second criterion, the ability to detect changes in synchrony, we focused on true dyads and examined which parameter configurations were most sensitive to detecting changes in synchrony between the baseline and the story telling conditions. A paired t-test was used as a measure of distance between mean synchrony levels during storytelling and baseline, with a positive t-statistic indicating higher synchrony during storytelling. To ensure robustness, the analysis was run twice: once comparing stories told by Participant 1 with the baseline measures, and once for stories told by Participant 2 (see Supplementary Materials for the reasoning behind the choice of these comparisons). Only the first three minutes of each story were used to equate story length, and each comparison included both a positive and a neutral story given that a preliminary analysis revealed no differences between story valences. The only difference was that in the first comparison, Participant 1 told the stories and in the second comparison, Participant 2 told the stories. Being Participant 1 or 2 was based on the participant number and therefore should not have had any systematic impact on the synchrony level between the two individuals.

## Results

### Synchrony versus pseudosynchrony

#### Heartrate

Multiple parameter configurations successfully differentiated true from surrogate dyads (Figure 4a), with the best discrimination (t = 28.32) observed for the smallest window size (4 sec) and a maximum lag of 7.5 sec (Figure 5a). When mapping the t-statistics distribution onto the parameter configuration space, a clear pattern emerged: smaller window sizes consistently resulted in larger t-statistics, while excessively large window sizes caused synchrony estimates to flip, with surrogate dyads showing higher synchrony than true dyads, even more so when paired with small maximum lags (dark blue coloring in Figure 5a). This pattern is evident by the gradual changes in coloring from blue to yellow in Figure 5a when moving down the y-axis (i.e., moving from large to small window sizes). The maximum lag was less influential but still important on differentiating between synchrony from pseudosynchrony, with optimal performance around twice the window size (4 sec). Increasing or decreasing the maximum lag reduced the sensitivity to distinguish between the true and surrogate dyads as indicated by less yellow colors when moving left or right on the x-axis in Figure 5a. The replication analysis on the data from the second baseline confirmed this pattern (maximum t = 35.23, window size = 4 sec, maximum lag = 9 sec), with the maximum lag differed slightly by 1.5 sec showing the highest t-statistic at 9 sec. In conclusion, given the aim of the study to verify whether synchrony evolved as a result of interpersonal processes during a conversation above and beyond the shared environment of two participant, the range of parameters able to detect that difference is rather wide. In general, we recommend using a small window size for heartrate synchrony. Regarding the maximum lag, the choice of parameters is less influential, however, we recommend using a maximum lag that is around twice the window size.

**Figure 4.**
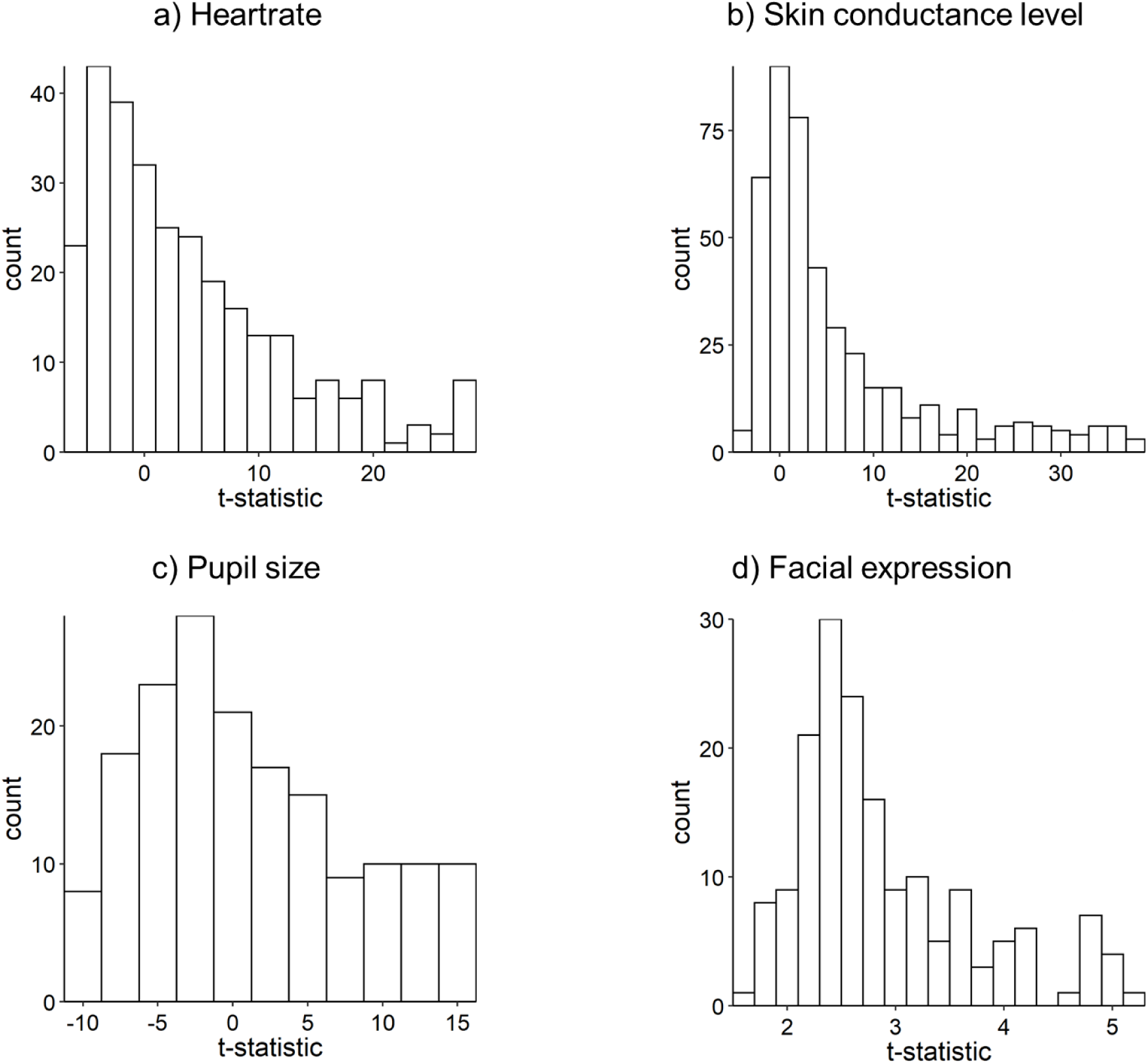
Distribution of t-statistics of the comparison between the true and surrogate dyads for each physiological measure. A positive value indicates higher synchrony level in the true compared to the surrogate dyads. Each data point represents one parameter configuration. For the analyses, data from the first baseline measure were used.

**Figure 5.**
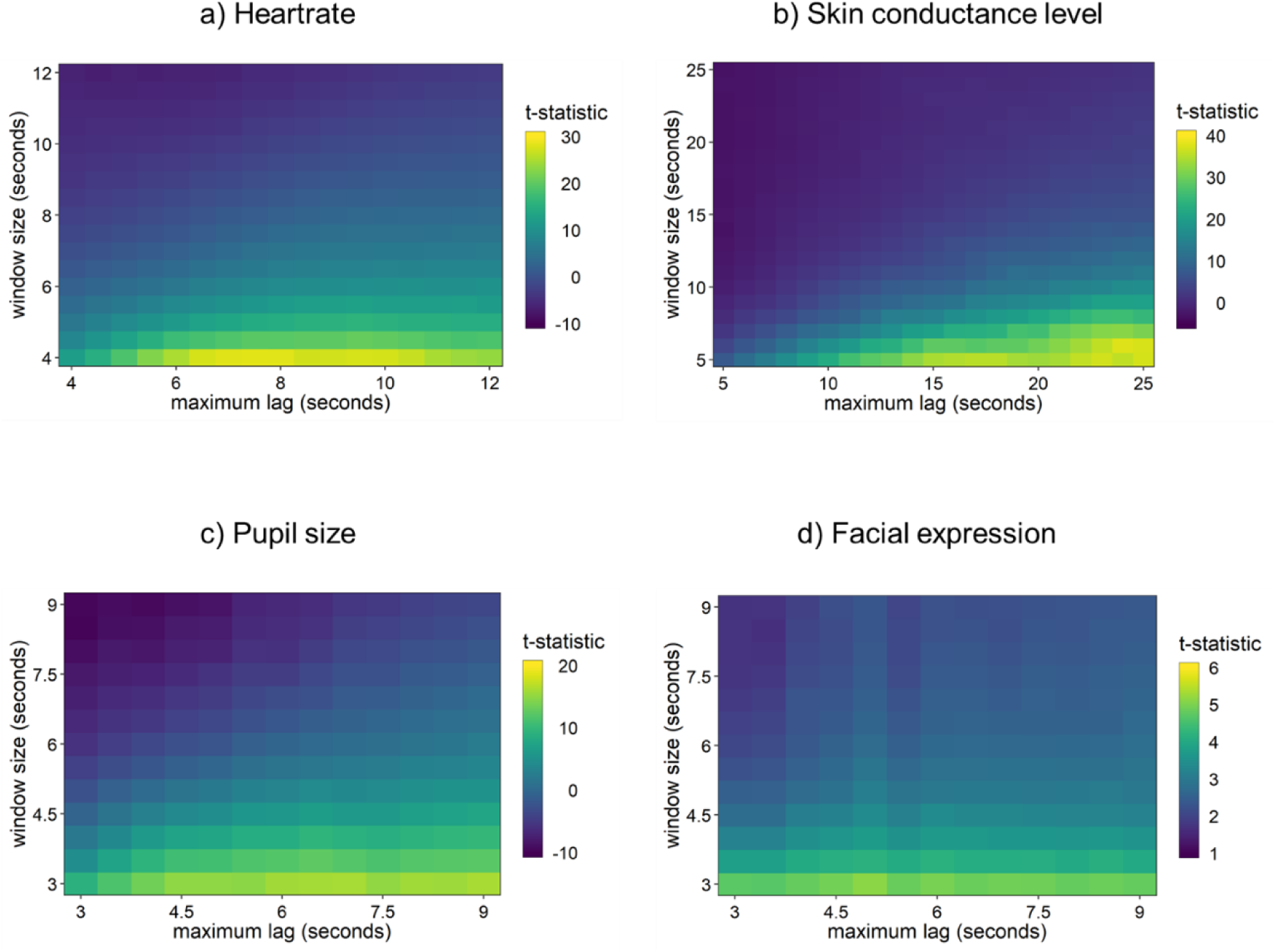
Distribution of the t-statistics of comparison between the originate and surrogate dyads for all parameter configurations and each physiological measure. The color coding runs from the lowest (blue) to the highest (yellow) t-statistic. A positive t-statistic indicates that the true dyads showed higher synchrony levels than the surrogate dyads. The more yellow, the better the discrimination between the true and surrogate dyads. Data from the first baseline measure were used. Notice that the scaling of the axes and the color coding are adjusted to each physiological measure to increase comparability between parameters.

#### Skin conductance level

As with heartrate synchrony, Multiple parameter configurations were sensitive to distinguish the true from the surrogate dyads (see Figure 4b). The largest t-statistic of 37.71 was observed for a window size of 6 sec and a maximum lag of 24 sec (see Figure 5b). Excessively large window sizes paired with small maximum lags caused synchrony estimates to invert, with surrogate dyads appearing more synchronized than true dyads. Notably, unlike heart rate, discriminative ability steadily increased when small window sizes were combined with progressively larger maximum lags (around four times the window size), a pattern replicated in the replication analysis (maximum t = 48.71; window size = 5 sec, maximum lag = 21 sec - see Figure S1b-S2b). Based on these results, we recommend using a small window size and a large maximum lag that is around four times the window size.

#### Pupil size

Also in this case, the number of positive t-statistics depicted in Figure 4c indicates that several parameter configurations could differentiate synchrony from pseudosynchrony. The maximum t-statistic of 16.12 was associated with a window size of 3 sec and a maximum lag of 9 sec. The general pattern as for the other measures was observed: the smaller the window size, the greater the difference between the true and surrogate dyads, while and excessively large window size would estimate larger synchrony for the surrogate compared to the true dyads (see Figure 5c). The maximum lag was less influential than the window size, showing only a slight tendency toward larger values. The replication analysis confirmed this pattern (maximum t = 18.04; window size = 3 sec, maximum lag = 6.5 sec). In conclusion, smaller window sizes were more sensitive to distinguishing synchrony from pseudosynchrony in pupil size data. The maximum lag did not have as much of an impact, but should be set to two to three times the window size.

#### Facial expression

All t-statistics were positive indicating that the level of synchrony was higher for the true compared to the surrogate dyads for all parameter configurations, though with less variance than the other measures (range: 1.68 to 5.14; Figure 5d). The best discrimination (t = 5.14) was observed for the smallest window size (3 sec) and a maximum lag of 5. The same pattern as for the other three measures emerged: the smaller the window size, the better the true dyads could be distinguished from the surrogate dyads. Maximum lag was not influential on the discriminative ability, but the largest t-statistic was observed at almost twice the window size (5 sec). For the replication analysis, a similar pattern was observed (maximum t = 7.05; window size = 3 sec, maximum lag = 8.5 sec) with a slightly wider range of t-statistics (range: - .16 to 7.05; Figure S1d-S2d). In conclusion, we recommend using a small window size and a maximum lag that is two to three times the window size.

### Change in synchrony

#### Heartrate

The largest absolute t-statistic was negative indicating that synchrony levels were higher during baseline compared to storytelling (see Figure 6a). The highest absolute t-statistic of 4.86 was observed when the window size was set to 4 sec. Similar to the first comparison analysis, smaller window sizes could discriminate the two conditions better than large window sizes (see Figure 7a). Again, the maximum lag was less influential than the window size parameter, but the best outcome was observed for the smallest maximum lag of 4 sec. The absolute t-statistic steadily decreased with increasing maximum lags. Replication analysis showed similar results, with smaller window sizes showing the greatest discriminative power between the conditions (see Figure S3a-S4a). Specifically, the largest absolute t-statistic was again observed for a window size of 4 sec, with only minor variation across maximum lags (optimal maximum lag = 7 sec). Therefore, based on both analyses the conclusion is: if the aim is to distinguish synchrony levels in heartrate responses between two (within-subject) conditions, the smaller the window size, the better. The maximum lag is less influential, but its optimal value should be equal to or twice the window size.

**Figure 6.**
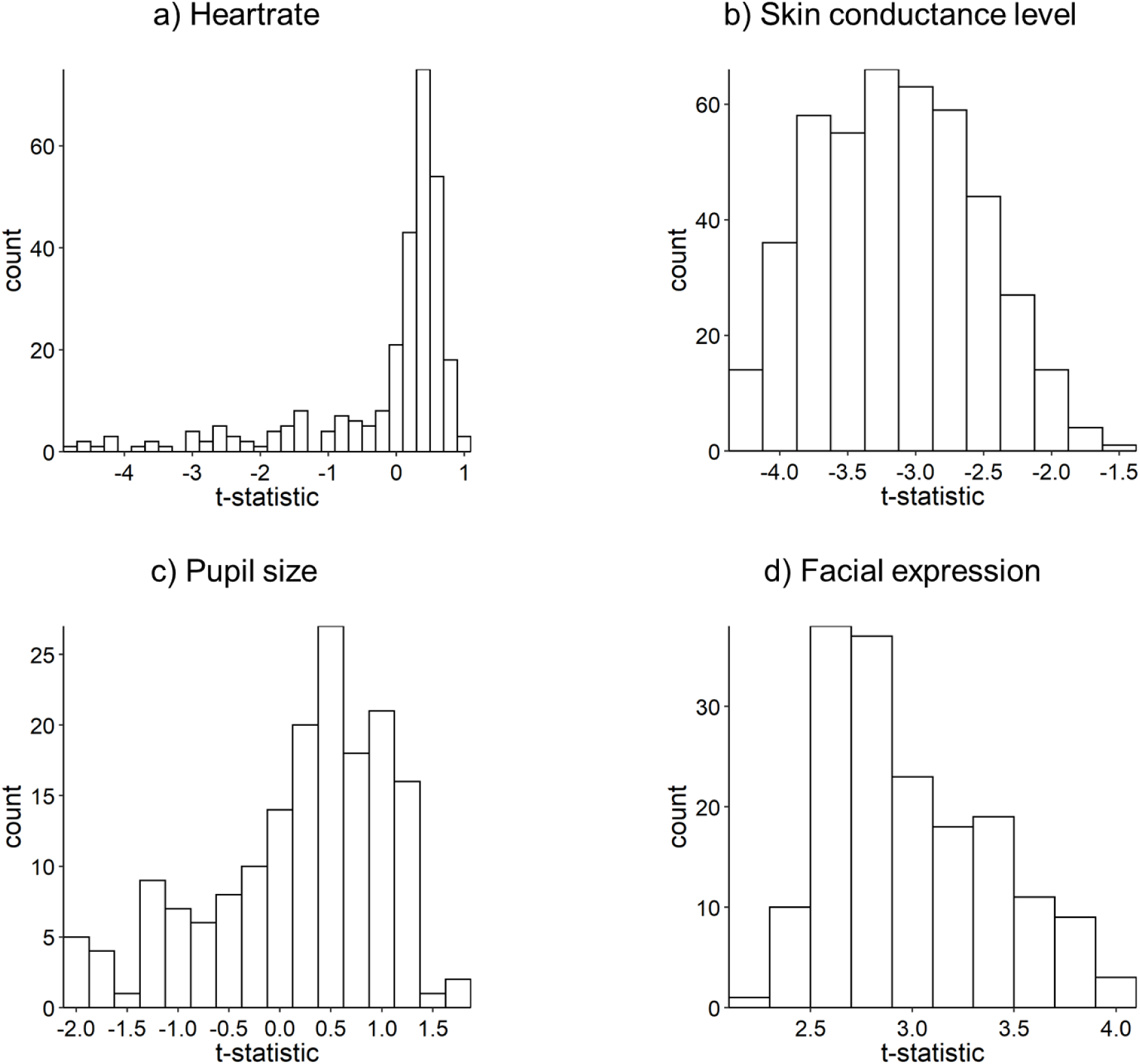
Distribution of t-statistics of the comparison between storytelling and baseline for each physiological measure. A positive value indicates higher synchrony levels during storytelling compared to baseline. Each data point represents one parameter configuration.

**Figure 7.**
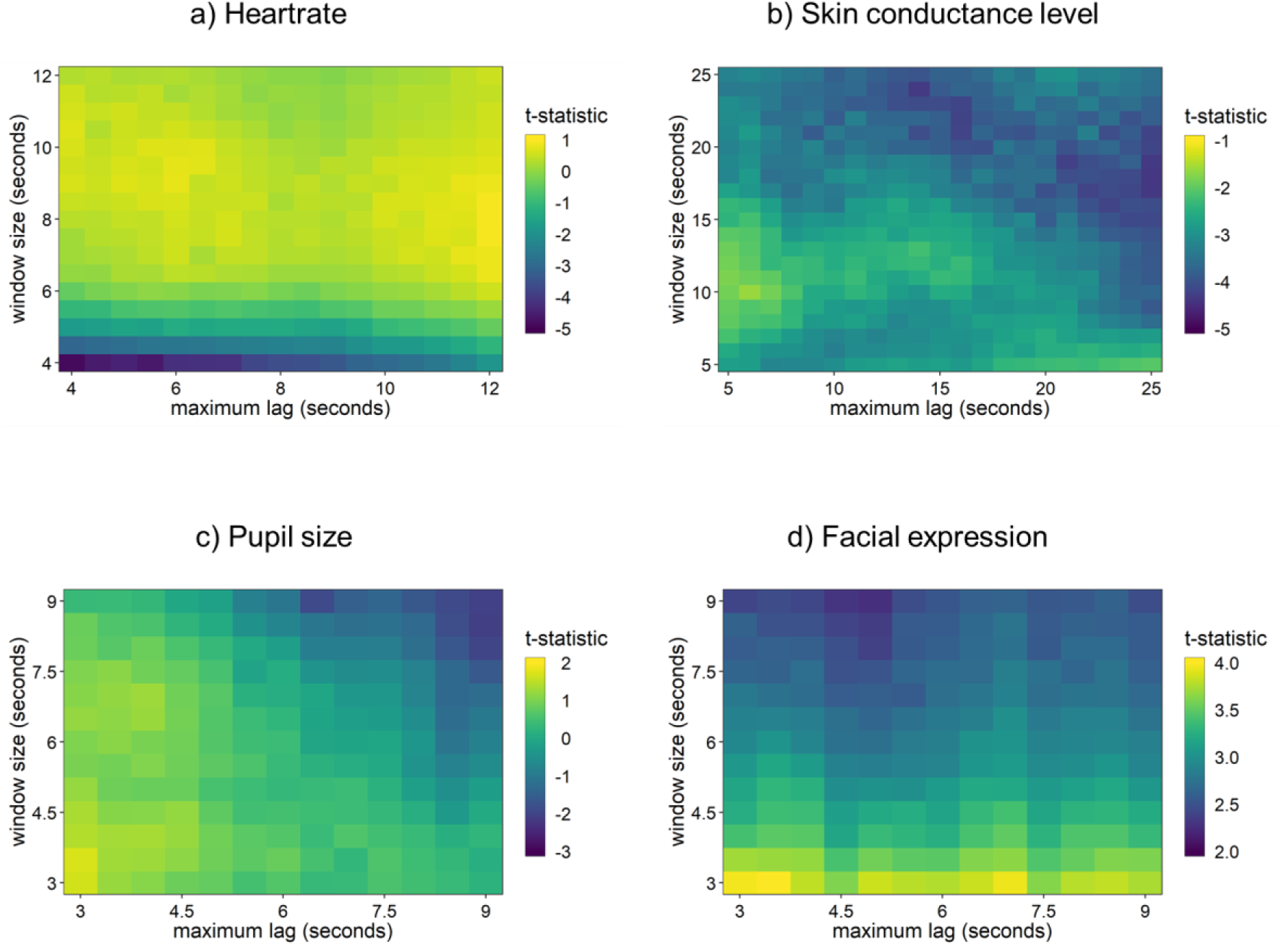
Distribution of t-statistics of the comparison between storytelling and baseline of all parameter configurations for each physiological measure. The color coding runs from the lowest (blue) to the highest (yellow) t-statistic. A positive t-statistic indicates that the level of synchrony was higher during storytelling than during baseline. Analysis was based on data from both baseline measures and the first and third stories. Notice that the scaling of the axes and the color coding are adjusted to each physiological measure to increase comparability between parameters. Also, the highest t-statistic was not always the highest absolute value with the latter value being discussed in the result section. However, the general idea of greater (absolute) t-statistics indicating better discrimination between the two conditions remains.

#### Skin conductance level

All t-statistics were negative indicating that the level of synchrony was higher during the baseline measures compared to during storytelling (see Figure 6b). The highest absolute t-statistic of 4.37 was observed for a window size of 18 sec and a maximum lag of 25 sec. Interestingly, the recurrent pattern of smaller window sizes showing greater t-statistics was not evident (see Figure 7b). Even though there seemed to be a weak tendency for absolute t-statistics to become larger with larger window sizes and maximum lags, the pattern was rather weak. In addition, the difference between t-statistics was small (range: - 1.61, -4.37). The replication analysis deviated substantially, with the largest absolute t-statistic observed for a window size of 5 sec and a maximum lag of 12 sec, and an even narrower range (-.19 to -2.56; Figure S3b-S4b). Given the lack of clear patterns in the parameter configuration space and considerable discrepancies in the results between the primary and replication analyses, we cannot draw meaningful conclusions about the optimal parameter configuration for detecting changes in skin conductance synchrony.

#### Pupil size

Parameter configurations strongly influenced the direction of results, with small window sizes and maximum lags yielding higher synchrony during storytelling, and large window sizes and maximum lags yielding higher synchrony during baseline (Figure 7c). The largest positive t-statistic (1.72; window size = 3.5 sec, maximum lag = 3 sec) and largest absolute t-statistic (2.07; window size = 8.5 sec, maximum lag = 9 sec) pointed in opposite directions, a pattern that was replicated with identical optimal parameter values. Given the ambiguity across parameters, we refrain from providing recommendations for this measure and instead caution that parameter choices can substantially alter the conclusions drawn about changes in pupil synchrony between conditions.

#### Facial expressions

All t-statistics were positive, indicating higher synchrony during storytelling than baseline (Figure 6d). The best discrimination was observed for the smallest window size (3 sec; t = 3.99), with a weak but consistent pattern of smaller window sizes yielding larger t-statistics (Figure 7d). Albeit weak, the general pattern emerged with larger t-statistics being associated with smaller windows sizes. The maximum lag had negligible influence on the results, with the biggest difference between conditions was 3.5 sec. The replication analysis confirmed this pattern (maximum t = 4.53; window size = 3 sec), with only minor variation across maximum lags (Figure S3d-S4d). To conclude, if the aim is to detect change in synchrony between two conditions in facial synchrony, then the window size should be set to a small value. The effect of the maximum lag was negligible, however, to be consistent with the other measures, we recommend a maximum lag twice the window size.

## Discussion

The fact that we synchronize with each other’s emotional expressions and physiological states has intrigued researchers in many different disciplines. The present study investigated optimal parameter configurations for Windowed Cross-Correlation (WCC) analysis applied to four physiological measures: heartrate, skin conductance level, pupil size, and activity of the *Zygomaticus Major* muscle (associated with smiling). We evaluated parameter configurations on two criteria: the ability to distinguish synchrony from pseudosynchrony, and the sensitivity to detect changes in synchrony between conditions.

Regarding the first criterion, a consistent pattern emerged across all four physiological measures: smaller window sizes more effectively discriminated true from surrogate dyads, namely synchrony from pseudosynchrony. With respect to the second parameter, the maximum lag was generally less influential and larger than the corresponding window size. Crucially, this pattern was not specific to any particular signal but may reflect an intrinsic characteristic of how cross-correlations are calculated, a point we elaborate on below.

Regarding the second criterion, results were less conclusive. Here, we compared baseline measures where people looked at each other in silent with periods where participants engaged in storytelling. While heartrate and facial expressions showed a weak tendency for smaller window sizes to better detect condition differences, no clear pattern emerged for skin conductance level, and pupil size results were highly sensitive to parameter choice, with small parameters favoring storytelling and large parameters favoring baseline. Two explanations are possible: (1) the difference between the two conditions was negligible and that the sensitivity to detect such small differences was barely affected by the parameters; (2) there were differences between the two conditions, but the method was not sensitive to detect them. In support of the first explanation, in two previous studies, we have used parameters included in the current analysis with which we were able to detect differences in within-subject conditions and could link it to interpersonal outcomes (Behrens et al., 2020; Prochazkova et al., 2019). The method therefore has been shown to be sensitive in other contexts. Yet, we refrain from address this using simulated data or more extreme experimental contrasts.

Table 2 summarizes our parameter recommendations. In what follows, we discuss our findings in depth and integrate them with theoretical considerations, focusing on the criterion to distinguish synchrony form pseudosynchrony.

**Table 2.** Summary of recommendations per parameter of the WCC analysis.

| Parameter | Recommendation |
| --- | --- |
| Window size | - Lower bound: large enough to capture meaningful information and variance within the signal of interest<br>- Upper bound: the response duration of the signal of interest; assumption of stationarity is met |
| Maximum lag | - Lower bound: at least as long as the window size<br>- Upper bound: at most twice as long as the window size |
| Window and lag increment | - Lower bound: 1 datapoint<br>- Upper bound: same as the window size / maximum lag<br>- Balance computational time and resolution: 1-5% of the window size / maximum lag |

### Window size

We observed that a wide range of window sizes were able to distinguish between synchrony and pseudo synchrony, with a recurrent pattern across measures: smaller window sizes generally performed better, while excessively large window size turned the table to the extent that the levels became lower for the true dyads than the surrogate dyads. To understand this general pattern, we should first quickly recap the purpose of the surrogate data analysis. The ultimate aim of this analysis is to destroy any synchrony that is the result of interpersonal processes while preserving all other statistical properties by generating new pseudo dyads that never actually interacted during the study. This way we knew that the null hypothesis, i.e., there is no interpersonal synchrony between participants, would be true; and we expected it to be true independently of the parameter configurations. Hence, the distribution of cross-correlations for the surrogate dyads stayed constant across all parameters. In contrast, for the true dyads, synchrony did emerge, which we expected based on prior research. During a dynamic interaction, however, there are moments when dyads are in sync, but also out of sync (Boker & Rotondo, 2002; Chen et al., 2020). These “anti-synchrony” moments are also referred to as antiphase synchrony or antiphase linkage (Reed et al., 2013), and they are the key element to explain the patterns we observed: if the window size is too large, both moments of synchrony and anti-phase synchrony are likely to be included into one window segment, averaging out and substantially reducing the strength of synchrony. This would result in synchrony estimates to invert, with surrogate dyads, likely lacking the synch-non-synch dynamics of a true interaction, appearing more synchronized than true dyads. Following the same logic, a smaller window size reduces the variance, causing the overall synchrony level to increase.

As seen in Equation 1, the cross-correlation is calculated by dividing the distance between each datapoint and the mean of the window segment by its standard deviation: the smaller the window size, the less chance for variation to occur within a window (i.e., the smaller the standard deviation), the higher the correlation. In this case, while the distribution of correlation estimates stayed constant for the surrogate dyads, the estimates for the true dyads increased, the distance between the means of these two groups became increasingly larger, and the general pattern we observed across measures emerged. Thus, this pattern is not measure-specific but rather reflects an intrinsic mathematical property of cross-correlation which applies to all time-series.

One may wonder whether steadily decreasing the window size will also steadily increase the ability to distinguish synchrony from pseudosynchrony. The short answer is no. Imagine the extreme case, where the window size consists of two data points. These two datapoints hold very little information and would only allow for possible correlation values of -1 and 1. This reduces the sensitivity for measuring synchrony and therefore for distinguishing synchrony from pseudosynchrony. Consequently, somewhere between a window size containing two datapoints and the smallest window sizes we examined, there will be a turning point, where the two types of dyads will become distinguishable. Although statistically possible, making the window size as small as possible (but above the turning point) is not advisable for two reasons: (1) a sufficient number of data points are needed to reliably estimate correlation coefficients (Schönbrodt & Perugini, 2013), and (2) the window should capture a meaningful response. As outlined earlier, in order to reliably estimate a correlation coefficient, 65 to 250 data points are necessary, depending on the strength of the correlation. With a sampling rate of 20Hz across all measures, we therefore used a window size of at least 3 seconds (60 datapoints). If researchers want to further decrease the window size, they should increase their sampling rate accordingly, whilst keeping in mind the biological plausibility of the signal investigated. With that said, a window size must include responses constrained by a meaningful upper and lower bound.

In the current study, we narrowed the possible values for the window size parameter by showing a range of parameters that qualify as potentially suitable. How can researchers choose between these options? To answer this question, we should go back to why cutting the time series into segments, i.e., reducing the non-stationarity in the signals, is relevant in the first place. A stationary signal has constant statistical properties, including, among others, a constant mean and standard deviation within that signal. In a dynamic interaction, the strength of synchrony will vary between moments of strong and weak synchrony. The window size needs to be small enough such that the synchrony level stays constant *within* that window. Determining how small the window must be, depends on the nature of the signal and is contained by an upper and lower bound. Let’s make some concrete examples. Imagine two interaction partners smiling at each other, with smiles coded in a binary fashion: a person either smiles or does not. Given an appropriate correlation measure for categorical variables, if the two participants smile at the same time for the same duration, there will be perfect synchrony between them for the entire duration of the two smiles. In this case, the window size could be as large as the duration of the smile because the level of synchrony is constant during that interval. However, if the smiling response occurs in a dynamic interaction and is continuously measured with the activity of the *zygomaticus major* (as in the current study), there will be variations in latency, magnitude, and duration of the smiles within and between individuals. In this case, the level of synchrony is likely to change even within the window that would have been categorized as a ‘smile’ in the discrete scenario described above: one person might show a long, pronounced smile, while the other might smile later and for a shorter length of time, and synchrony would only occur during the short time when both people simultaneously smile. Therefore, if a smile is detected through facial muscle activity, the window size should be smaller than the duration of a “typical” smile to capture these variations, optimally at most half the response duration. In this way, at least two estimates of the level of synchrony will be computed for that response, capturing changes in synchrony that are twice the speed of the overall response. Other than the upper bound for window size being smaller than the response duration of interest, there is a lower bound as well. In particular, the window size should be large enough to capture meaningful variations within a response. For example, if the signal of interest is skin conductance level, a window size of 1 second would contain straight lines in most windows. This would produce extreme cross-correlations without capturing meaningful changes in the signal. On the other hand, applying the same window size to facial expressions might be considered a medium to large window, given that a smile has been shown to last 500ms to 4 seconds (Frank et al., 1993). Both the upper and lower bound therefore, determine the potential values for the window size.

We realize that defining the “duration of a response” in physiological signals is inherently difficult due to substantial variability within and between individuals. It is beyond the scope of the current paper to provide concrete guidelines for this matter; researchers must decide which aspects of the signal are theoretically relevant for their research question and design. The appropriate (range of) window size(s) likely differ across situations and conditions, and should be treated as a testable assumption, namely, that responses synchronize that are equal to or longer than the window size chosen. Faster fluctuations may still be present but are likely attenuated through averaging.

In the absence of strong a priori hypotheses, researchers may examine a limited range of window sizes to determine which timescales capture synchrony in a given context. Comparing two to three potential values can shed light on the temporal scale at which synchrony occurs in a particular context. Of course, because synchrony is unlikely to occur only at a single fixed duration, similar results are to be expected for window sizes closer together. Consequently, referring, for instance, to “skin conductance synchrony” based on a single parameter choice risks overgeneralization. Our recommendation is to preregister the parameter range one intends to use and to report the distribution of correlation coefficients across these parameters. This approach would make the parameter choice transparent while strengthening the interpretability of conclusions.

To conclude, the results of the current study indicate that a range of window sizes are able to detect synchrony that occurs as a result of interpersonal processes with a preference for shorter window sizes. From a theoretical perspective, the range of potential window sizes is contained by (i) an upper bound defined by the length of the duration of the responses under investigation and (ii) a lower bound defined by sufficient variation within the window. Rather than searching for that one most suitable parameter for each physiological measure, choosing a window size should be seen as a hypothesis being tested. Importantly, researchers need to be specific about what aspect of a signal they investigate, which should be clearly stated in both their hypothesis and conclusions.

### Maximum lag and increments

Our results revealed that the maximum lag was less influential than the window size, yet not trivial. In the Results section, we included recommendations on how to specify this parameter in relation to the window size. For three of the four measures, the optimal maximum lag was around twice the corresponding window size. For skin conductance level, this ratio increased to four times the window size. However, we would like to point out that the corresponding window size of 4 sec was very small. Considering the slow time course of the measure, a larger window size is likely to be preferred, and we expect the ratio to converge with the one recommended for the other measures. In Table 2, we set a maximum lag twice the window size as the upper limit, which means that there can be up to a full response between the responses of the interacting individuals. For example, imagine the measure of interest is facial activity, and the window size is 2 seconds: if the maximum lag is 4 seconds, then two smiles that occur 4 seconds apart from each other are considered synchronized responses. This situation seems still reasonable in the context of a real conversation, yet on the upper limit. However, expanding the maximum lag to 6 seconds (a maximum lag three times the window size) would likely link two unrelated events to one another. The lower bound of setting the maximum lag equal to the window size corresponds to the situation where people respond to each other in direct succession. In sum, as a general rule of thumb, we recommend a maximum lag of at least equal to and at most twice the size of the window size. We adjusted the increments such that the windows and lags moved by 10% of the window size and maximum lag, respectively. This was a choice of practicality, reducing the computational time in light of the huge number of analyses run while keeping the resolution sufficiently high. If researchers focus on specific parameters, we speculate that reducing the percentage to 1 to 5% offers a good balance between analysis sensitivity and computational time. However, future research is needed to determine the optimal balance between sensitivity and computational time.

### Limitations

There are a few limitations that we would like to point out. First, in the current study, we concentrated on the window size and maximum lag parameters, while setting the window and lag increments to 10%. A systematic investigation of the effect of changing these parameters is needed. As mentioned earlier, estimations of the level of synchrony will stabilize with smaller increments, such that decreasing the increments even further will add little information at the cost of extra computational time. Second, the guidelines we propose in Table 2 may not be generalizable to other measures of synchrony and may not be applicable to other biological time series. Researchers should therefore be careful with making any inferences about other statistical analyses and time series other than those used in the current study. Third, all data come from a single study and are subject to method variation. To reduce such variation, we ran all analyses twice with different data from the same study. However, this does not address method variations that are the result of the study itself, and future studies should replicate our findings in a different dataset. Furthermore, we changed the original plan for the comparison of time intervals as outlined in “Choice of comparisons” in the Supplementary Materials. A more tailored study design may have found more specific results, in particular with regard to the sensitivity to detect change in synchrony. Related to that, WCC analysis only detects moments where the responses of two individuals move in the same direction (e.g., HR of both participants increases). Given the turn-taking nature of a conversation (i.e., one person talks while the other is listening), responses moving in opposite directions (e.g., antiphase synchrony) can also give rise to valuable information of an interaction as discussed by Reed et al. (2013) and Chen et al. (2020). This form of synchrony could be potentially captured with the WCC analysis by detecting the maximum correlation based on the *absolute* values rather than only looking at the positive values. However, we do not recommend making this change without considerable validation first. Finally, WCC captures temporary *linear* relationships between two time series. Therefore, non-linear patterns or other causal effects not captured by a correlation are not detected by the current method.

### Future directions and conclusions

The most important lesson the current study teaches us is that researchers need to be more precise in what they (aim to) investigate as defined by the parameters specified in the analyses. In the current study, dyads synchronized on a range of response windows. However, this might not always be true, especially if the aim is to link it to specific psychological processes that might be influenced by only particular physiological processes. Future studies should therefore make more refined distinctions of which components of a particular physiological signal are involved in the process of interest and how the different components interact. This will facilitate making well-informed decisions about the response windows and shed more light on the biological underpinnings of psychological processes.

Before making well-informed decisions on the parameter configurations *within* a particular method, it is important to realize what the differences are *between* methods. WCC analysis is one of many possible methods, and each method has its strengths and weaknesses. While one method might be appropriate for an experiment, it might not be for another, depending on, among others, the type of data (e.g., continuous or categorical measures), the measure of interest (e.g., strengths versus frequency of synchrony; global versus time-sensitive measure of synchrony) (Gates & Liu, 2016; Schoenherr et al., 2018), and the specific tasks involved. For example, we chose to treat facial muscle activity as a continuous measure, but researchers might also be interested in investigating discrete events of, for example, smiling and its synchrony in a conversation. In this case, the analysis developed by Altmann (2011), where time series are first categorized into intervals of synchrony and intervals of no synchrony before measures of the strength and frequency of synchrony are computed, may be more appropriate. In addition, even when using the same data, the outcomes can be somewhat different: Schoenherr et al., (2018) performed an exploratory factor analysis on seven linear time series analyses and different outcome variables, including the WCC analysis. They reported that all these methods measure the overall phenomenon of synchrony, but could be categorized into three correlated, yet distinct facets of synchrony: the strength of synchrony of the total interaction, the strength of synchrony during synchronization intervals, and the frequency of synchrony (Schoenherr et al., 2018). The WCC analysis as performed in the current study reflected the first facet. Researchers should therefore refine which facet of synchrony they are interested in and choose the appropriate methods accordingly.

The aim of the current study was to optimize the parameters for the WCC analysis from a statistical point of view. The initial idea was to provide researchers with concrete guidelines on which specific parameters would be most appropriate for the four physiological measures. However, the results show that when the aim is to detect synchrony, the parameters follow a general pattern that is not specific to the signal of interest, but rather a result of intrinsic characteristics of how the cross-correlation is calculated. That does not mean that the parameters should not be tailored to the signal of interest. Instead, theoretical considerations should be integrated with the findings observed in the current study. Here, there is no one-fits-all solution, which might not be surprising given that we aim to capture a highly complex dynamical process. The current study narrows down the range of possible parameters, and we provide guidelines on how to tailor the parameters further to the interests of the researcher. Being specific and transparent about these choices will increase the comparability across studies and add more and more pieces to the puzzle of understanding the phenomenon of synchrony.

## Acknowledgments

We thank the students who have assisted in collecting the data for this study. The work has been previously shared as a preprint:

Behrens, Moulder, Boker & Kret (2020) - Quantifying Physiological Synchrony through Windowed Cross-Correlation Analysis: Statistical and Theoretical Considerations at this link https://doi.org/10.1101/2020.08.27.269746.

## Author contributions

Conceptualization: F.B., R.G.M, M.E.K.; Data curation: F.B.; Formal analysis: F.B., R.G.M.; Investigation: F.B., R.G.M.; Methodology: F.B., R.G.M, M.E.K.; Supervision: M.E.K, S.M.B; Visualization: F.B., R.G.M.; Writing – original draft: F.B.; Writing – review & editing: R.G.M., F.D., S.M.B., M.E.K.

## Data and materials availability

All data, code, and materials that are associated with this paper and used to conduct the analyses are available on the Open Science Framework at this link: https://osf.io/vxm5b/overview?view_only=93ac3e21966642968b260dcbc9c061f6

