## Supplementary materials for "Quantifying Physiological Synchrony Using Windowed Cross-Correlation Analysis: Statistical and Theoretical Considerations"

**Choice of comparisons**

The initial plan was to make two comparisons for the "change in synchrony" criterion: i) compare the first baseline measure with the breathing exercise interval, and ii) compare the positive and neutral stories. Regarding the first comparison, the breathing exercise was meant to manipulate synchrony as people were explicitly instructed to breathe synchronously. Although this manipulation worked for the heartrate measure, there were no differences between baseline and breathing intervals evident in the other three signals across parameter configurations. Similarly, based on a previous study showing more synchrony during emotional periods during storytelling, we expected differences in synchrony between the positive and neutral stories in the current study (Kang & Wheatley, 2017). However, across parameter configurations the distance in means between the two conditions were negligible for all four physiological measures, despite significant differences in ratings for both the valence and the intensity ($M_{pos}$ = 7.53, $M_{neu}$ = 5.65, $t(135) = 15.49$, $p < .001$; $M_{pos}$ = 5.32, $M_{neu}$ = 3.13, $t(135) = 10.31$, $p < .001$, respectively). We could have used the comparison between the baseline measure and the breathing exercise for the heartrate measure, however, we wanted to use the same conditions across signals to be consistent between measures. Additionally, we wanted to prevent losing collected data and use comparisons that would be comparable for the primary and replication analysis. We therefore decided to include two intervals per condition and compare the two baseline measures with two storytelling intervals. It is important to note that the aim of our study was not to find differences between the conditions to support a theoretical research hypothesis. Instead, we wanted to perform comparisons between conditions where the difference was as large as possible and where variance between parameter configurations could potentially show. Using data without any effects observed across parameter configurations would raise the question of whether the results were due to a lack of actual differences or an insensitive method. Unfortunately, we still faced exactly that dilemma in the results. Nevertheless, we think that the chosen comparisons had the most potential to show effects and possible differences in parameter configurations.

Table S1

*Descriptive statistics of the Positive And Negative Affect Schedule (PANAS), the story ratings, the Interpersonal Reactivity Index (IRI), and the Five Facet Mindfulness Questionnaire (FFMQ)*

| **Questionnaire** | **Mean** | **SD** | **Minimum** | **Maximum** |
|---|---|---|---|---|
| *PANAS positive scale:* | | | | |
| Baseline | 2.69 | .70 | 1.20 | 4.60 |
| Positive stories | 2.75 | .74 | 1.00 | 4.40 |
| Neutral stories | 2.38 | .76 | 1.10 | 4.10 |
| *PANAS negative scale:* | | | | |
| Baseline | 1.56 | .44 | 1.00 | 2.70 |
| Positive stories | 1.29 | .38 | 1.00 | 2.50 |
| Neutral stories | 1.22 | .29 | 1.00 | 2.40 |
| *Story rating (valence):* | | | | |
| Positive stories | 7.54 | .91 | 5 | 9 |
| Neutral stories | 5.74 | .97 | 3 | 8 |
| *Story rating (intensity):* | | | | |
| Positive stories | 5.46 | 1.90 | 1 | 9 |
| Neutral stories | 3.23 | 1.74 | 1 | 7 |
| *Questionnaires:* | | | | |
| IRI | 3.16 | .25 | 2.47 | 3.82 |
| FFMQ | 2.99 | .25 | 2.25 | 3.58 |

*Note.* SD = standard deviation.

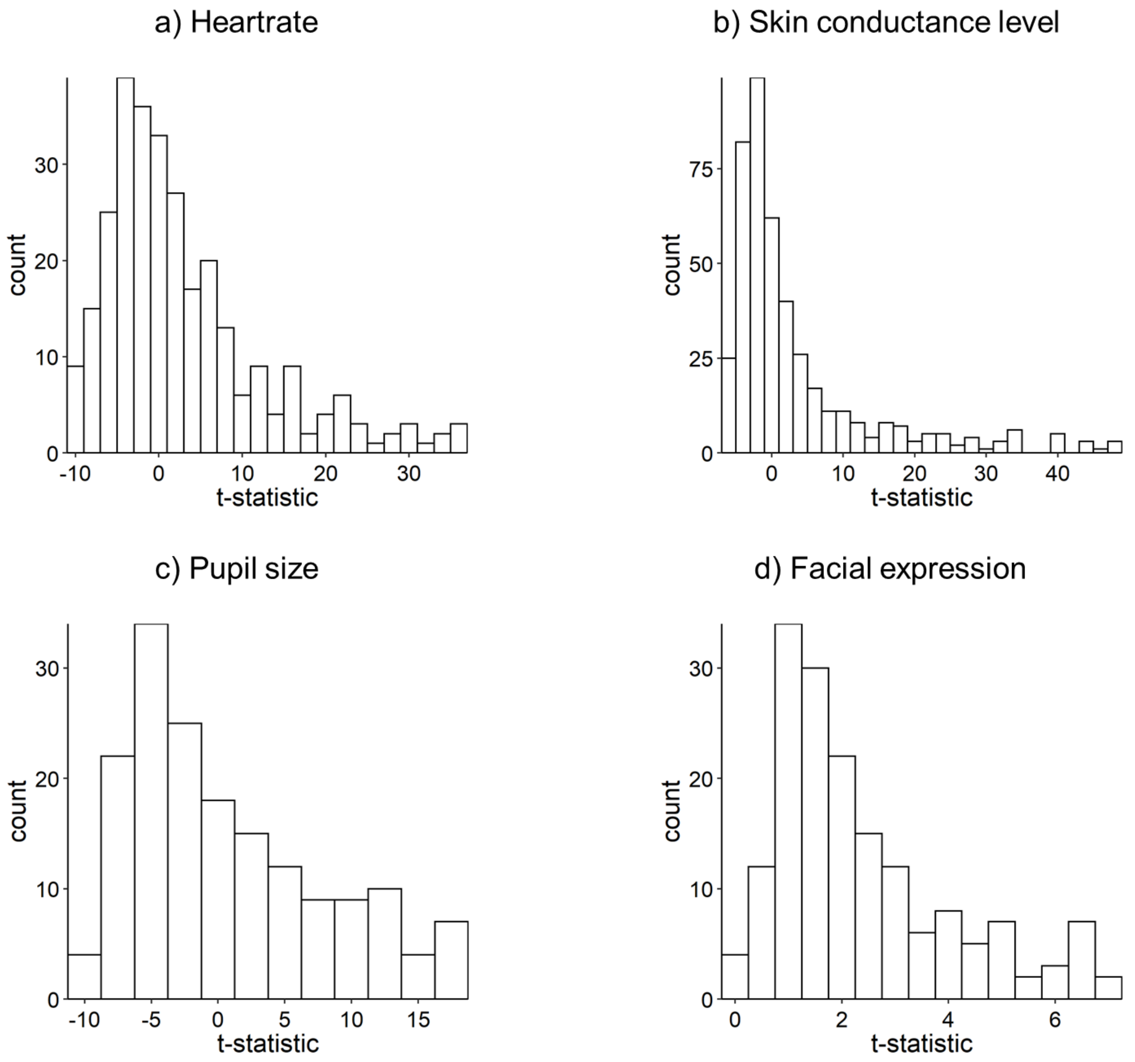


*Figure S1.* Distribution of t-statistics of the comparison between the true and surrogate dyads for each physiological measure (replication analysis). A positive value indicates higher synchrony levels in the true compared to the surrogate dyads. Each data point represents one parameter configuration. For the analyses, data from the second baseline measure were used.

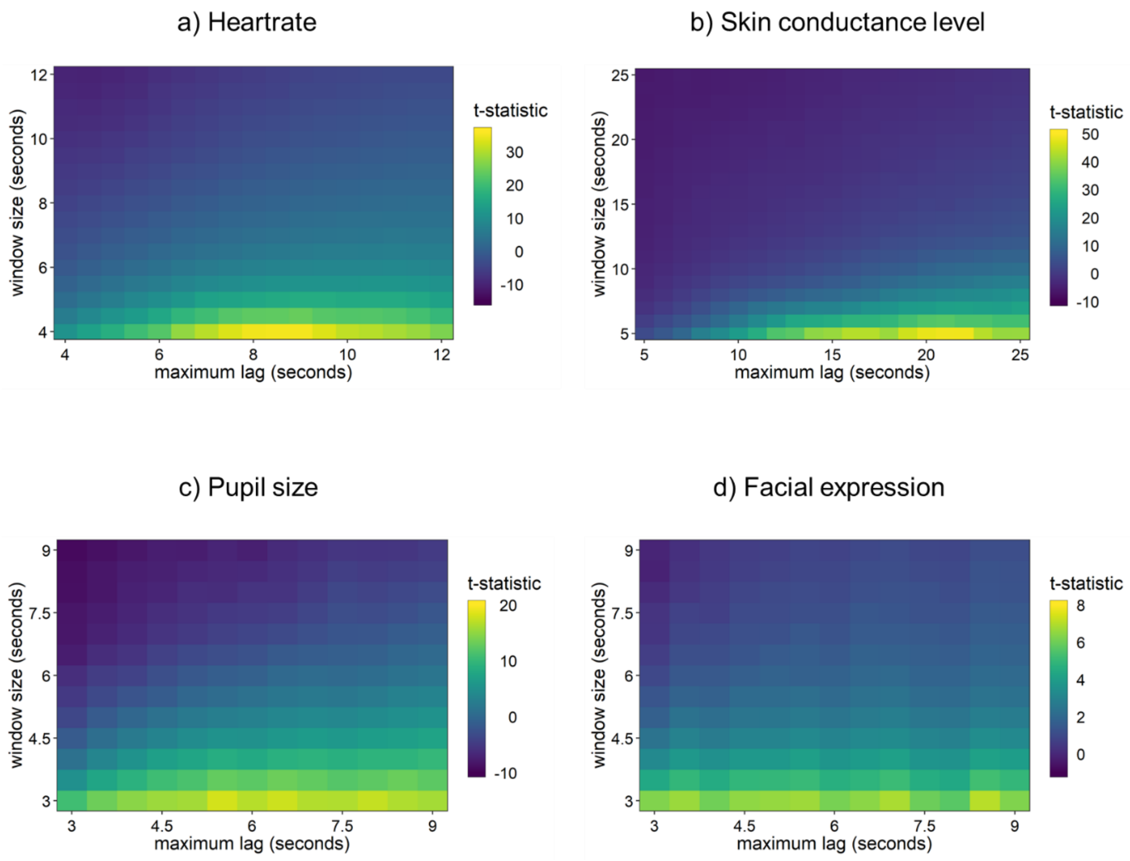


*Figure S2.* Distribution of the t-statistics of the comparison between the true and surrogate dyads for all parameter configurations and each physiological measure (replication analysis). The color coding runs from the lowest (blue) to the highest (yellow) t-statistic. A positive t-statistic indicates that the true dyads showed higher synchrony levels than the surrogate dyads. The more yellow, the better the discrimination between the true and surrogate dyads. Data from the second baseline measure were used. Notice that the scaling of the axes and the color coding are adjusted to each physiological measure to increase comparability between parameters.

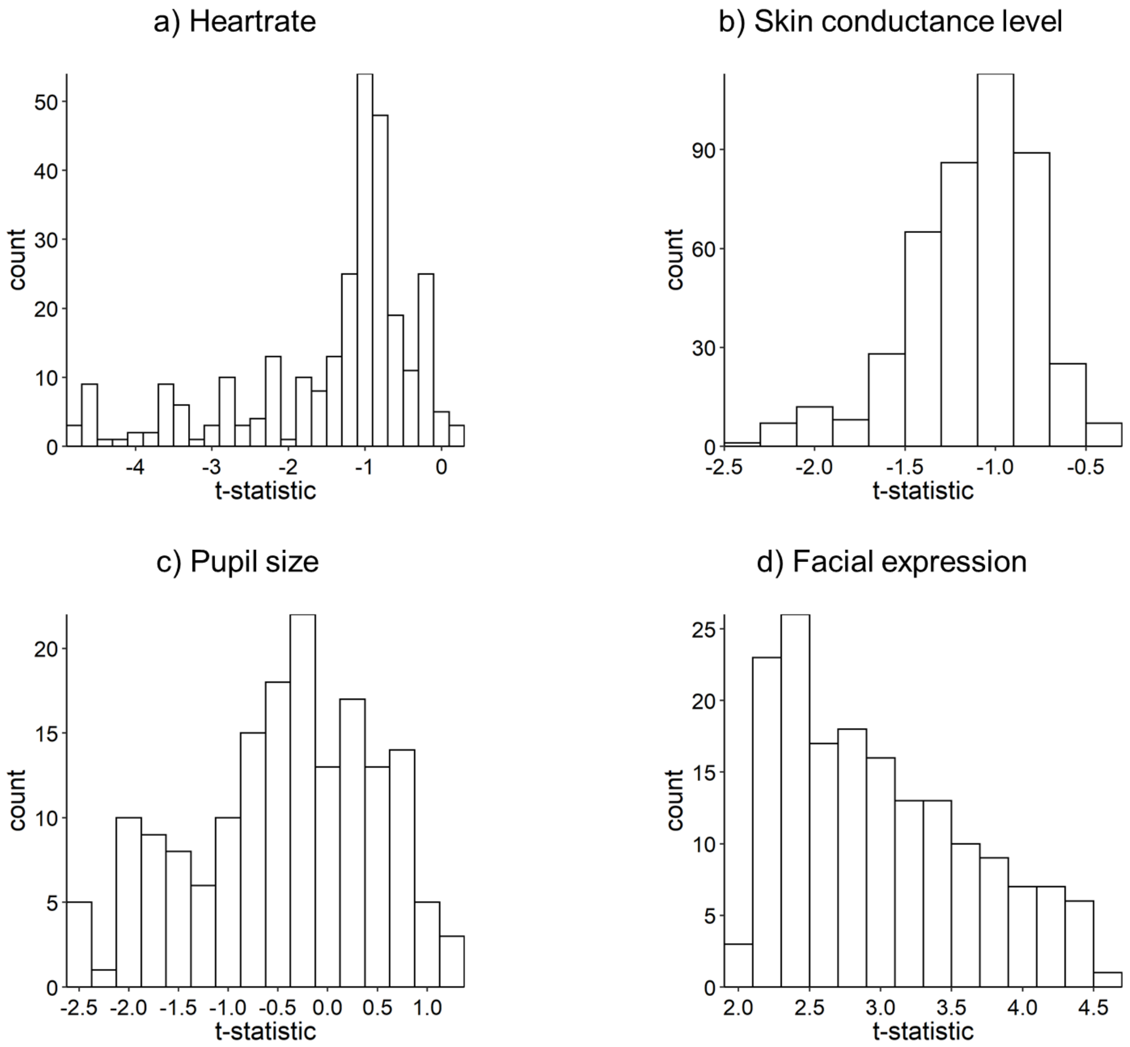


*Figure S3.* Distribution of t-statistics of the comparison between storytelling and baseline for each physiological measure (replication analysis). A positive value indicates higher synchrony levels during storytelling compared to baseline. Each data point represents one parameter configuration. Analysis was based on data from the second and fourth stories and both baseline measures.

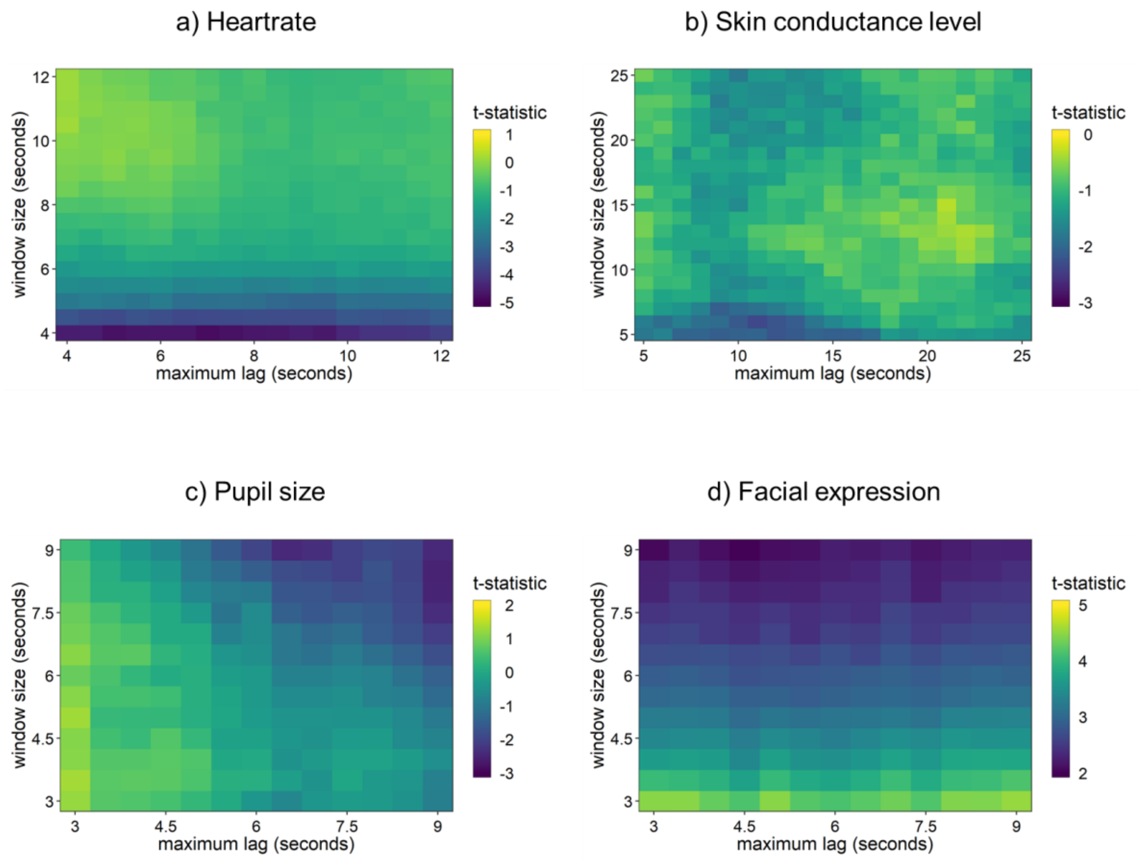


*Figure S4.* Distribution of t-statistics of the comparison between storytelling and baseline of all parameter configurations for each physiological measure (replication analysis). The color coding runs from the lowest (blue) to the highest (yellow) t-statistic. A positive t-statistic indicates that the level of synchrony was higher during storytelling than during baseline. Analysis was based on data from both baseline measures and the second and fourth stories. Notice that the scaling of the axes and the color coding are adjusted to each physiological measure to increase comparability between parameters. Also, the highest t-statistic was not always the highest absolute value with the latter value being discussed in the result section. However, the general idea of greater (absolute) t-statistics indicating better discrimination between the two conditions remains.
